# Probabilistic species distributions from nine large-scale gridded atlases over five decades

**DOI:** 10.64898/2026.09.24.754027

**Authors:** Florencia Grattarola, Gabriel Ortega-Solís, Carmen Diana Soria, Melanie Tietje, Friederike Johanna Rosa Wölke, Richard Fox, Lisbeth Hordley, Sergi Herrando, Petr Keil

## Abstract

Information on the long-term dynamics of species distributions is essential for assessing global biodiversity change and its causes, and for informed conservation decisions. A valuable source of such historical information are atlases. They provide spatial standardisation, extensive geographic coverage, temporal replication, documented sampling protocols, and span multiple species. Here we present model-based occupancy probabilities for 1,055 bird and 58 butterfly species, derived from nine gridded atlas datasets spanning several editions over the past 50 years. These are: Japan, New Zealand, New York state (US), the states of Alberta, Quebec, and The Maritimes (Canada), Czech Republic, United Kingdom, and Europe. Despite standardisation and coordination during data collection, the raw atlas data still contain some variation in sampling effort. We accounted for this using a two-step process: (1) we estimated a proxy for sampling effort in each grid cell within each atlas period using the Frescalo Algorithm, and (2) we ran a Bayesian spatial occupancy model for each species and atlas period, using the estimated sampling effort as the single predictor of detection in the observation process of the model. As the main product, we provide occupancy probabilities for species across grid cells and atlas periods. We also report uncertainty around each probability estimate and flag ∼21% (rare) species per atlas for which the model did not fit well. The probability distributions are fit for purpose and served in a user-friendly database with standardised geographic projection, metadata, and taxonomy.

## Background & Summary

The distribution of species across time and space is one of the core pieces of information in biodiversity science ^1^. Access to these data in an open digital form is more important than ever. Thus, it has been identified as one of the 2030 targets of the Kunming-Montreal Global Biodiversity Framework ^2^; Target 21: to improve accessibility to relevant biodiversity data, information and knowledge. A particular type of these data are atlases ^3–5^, which record species presences across contiguous grids, often temporally replicated over time in what are called atlas editions ^6,7^. Atlases are typically compiled at national or state levels and are coordinated by national government agencies, research institutions or not-for-profit organisations. To create them, the survey area is divided into grid cells (e.g., 10 x 10 km) and (mostly volunteer) surveyors visit these cells and record species following a set of recording guidelines. For example, participants might be asked to “complete two seasonal visits on three randomly selected squares and collect occurrence data in 10-minute intervals for a total of one hour” ^8^. In other cases, recording effort is left to the discretion of individual surveyors, but standardisation is maintained through the types of data collected and guidance encouraging complete recording of all species present. Atlas projects typically include rigorous data verification steps to ensure the accuracy of the generated data. The outcome is a list of expert-validated species presences per grid cell, collected over a period of time (e.g., four years) and, in many cases, repeated periodically using the same basic protocols.

Atlases are, however, not exempt from variation in recording effort ^9,10^. For example, a species can be missed in some grid cells because of relatively low recording effort, variation in recording strategies, or in recorders’ expertise. In some cases, there is also a lack of metadata on survey methods or the number of visits, making it difficult to account for the variation in effort in analyses. As a result, both spatial patterns ^11,12^ and temporal trends ^13^ derived from the raw atlas data can be inaccurate. To address this issue, several strategies have been proposed, such as thinning the data ^14^, or including sampling effort as a covariate in species distribution models (SDM) ^15,16^. The latter has been implemented in a category of SDMs called *hierarchical occupancy models* ^17,18^, which model the imperfect observation (detection) process jointly with the true distribution of the species. These models then predict the *probability of species occurrence or occupancy* (presence).

The estimated probabilistic distribution for each species in each region and time period can then be used in both basic and applied biodiversity research. For one, they represent an Essential Biodiversity Variable (EBV) ^1,19^, i.e. one of the core types of ecological information which directly underpins a range of biodiversity indicators. The probability distributions can be used, for example, to monitor temporal trends in species occupancy ^20,21^, or to test ecological hypotheses about the drivers of these changes ^22^. They can likewise serve as empirical benchmarks for mechanistic models of biodiversity ^23^, or as the basis for informed large-scale biodiversity assessments ^24^. Finally, the probabilities can also be combined with functional traits or phylogenetic information to study spatial patterns and temporal trends relevant to ecosystem functioning and to test evolutionary hypotheses ^25,26^.

Here we present model-based occupancy probabilities for 1,055 bird and 58 butterfly species derived from 9 gridded atlas datasets – from Japan, New Zealand, New York state (US), the states of Alberta, Quebec, and The Maritimes (Canada), Czech Republic, United Kingdom, and Europe, that span several editions over the past 50 years (Table 1, Figure 1). We derived these probabilities using two recent major advances, specifically the realisation that sampling effort can be approximated by local frequency of common (benchmark) species ^27^, and the emergence of fast and user-friendly implementation of the Bayesian occupancy models ^28^, which can combine the sampling effort with species detections to predict the probabilistic maps. The probabilities are presented in a common standardised format with resolved taxonomy, accompanied by full grid polygon files. This substantially reduces the cost of data management and cleaning, particularly when combining maps from multiple atlases.

**Table 1.**
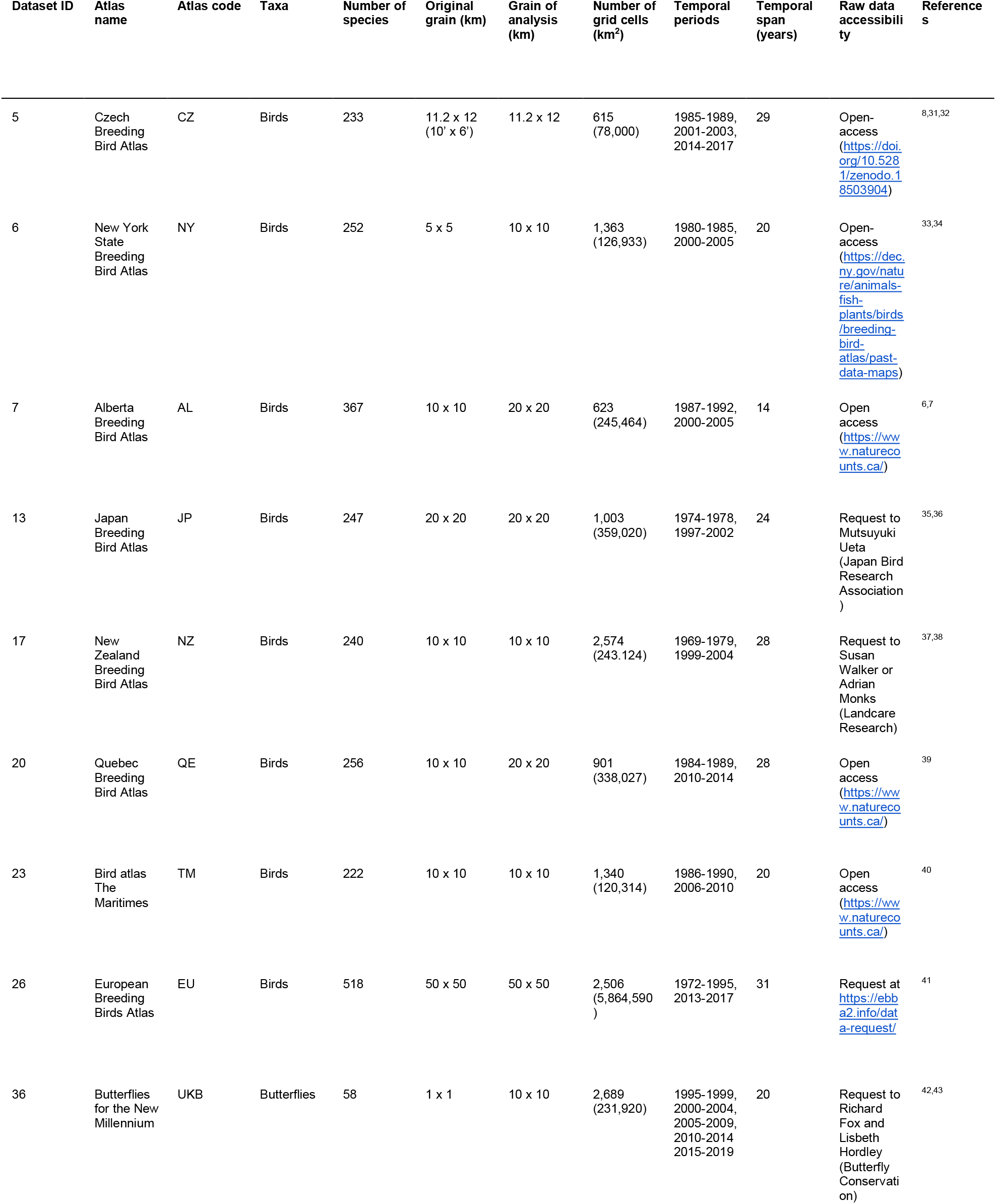
Temporally replicated gridded atlases for which sampling effort and species occupancy probabilities were estimated. The Dataset ID corresponds to the number in the internal Modelling of Biodiversity (MOBI) Lab database and was used for the file names.

**Figure 1.**
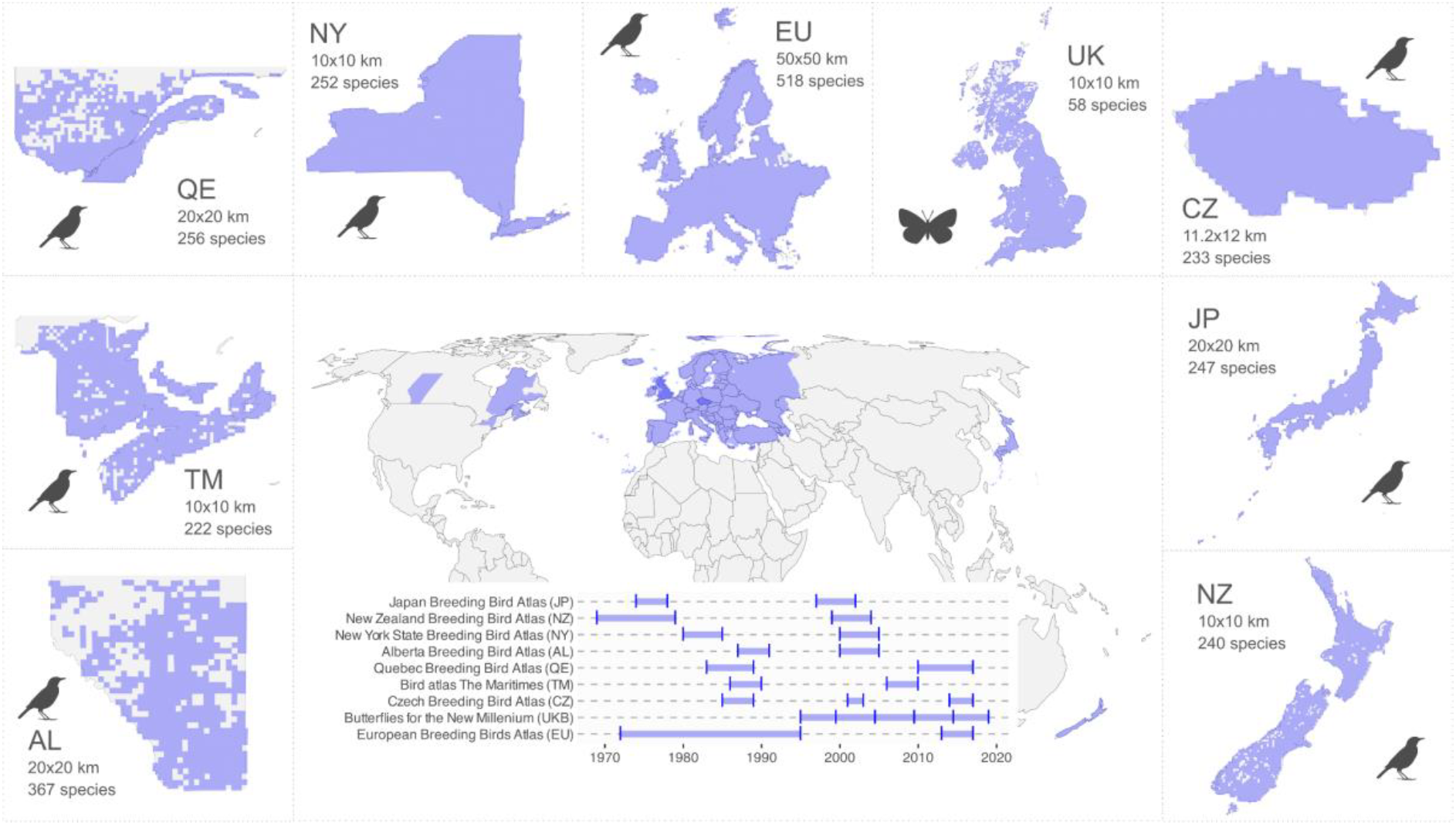
Temporal and spatial coverage of the atlases used for which we provide the probabilistic distributions. Each dataset spans a period between 14 and 31 years, over an extent of between 78,000 and 5,864,590 km2 (Table 1). Areas in darker blue (the Czech Republic and UK) are part of two different atlases, one at the continental and one at the national level.

The resulting dataset is fit-for-purpose and ready for immediate use ^29^: it is open access (CC BY), available for direct download in machine-readable formats, temporally replicated, and spatially comprehensive, with explicit spatial resolution, standardised taxonomy, and detailed metadata. Together, these properties make the dataset a uniquely interoperable resource for large-scale biodiversity research.

## Methods

### Atlas data

We obtained raw data from eight breeding bird atlases located in temperate regions (Japan, New Zealand, New York, Alberta, Quebec, The Maritimes, Czech Republic, and Europe), and one butterfly atlas (United Kingdom) (Table 1). We considered species detections as presences and assumed non-detections as absences. Some of the (original) datasets are openly available; others should be requested to the data owners. We provide this information below and in Table 1.

### Data characteristics

All atlas data consist of surveys conducted across standard grid cells that cover most of the country or state. The surveys were carried out by volunteers and professionals, following a specific protocol for each atlas (see below). Each sampling period within an atlas spans multiple years with repeated visits to most cells (Table 1). All bird species’ original names were harmonised according to HBW/BirdLife Taxonomic Checklist 9 (HBW & BirdLife International, 2024), while for butterfly species we retained the taxonomy used in the atlas, which follows Agassiz at el. ^30^. It should be noted, however, that taxonomy is far from constant over time and species splits can make comparisons between atlas periods extremely difficult. Although this type of taxonomic consideration should be carefully addressed when using repeated atlas work, it usually affects only a small minority of species. We recommend checking the species accounts in the original atlas works for further details.

Even though gridded atlases represent high-quality biodiversity data, there is inherent variation in field methods and the level of survey standardisation achievable, depending on the taxa studied and the geographic location of the grid cells. Here we present the main particularities of each dataset.

- Japan: It combines fieldwork records and questionnaire survey results. The fieldwork was done by 1 or 2 people walking along a pre-established survey route of ∼3 km and performing 2 x 30-minute point counts. This information was complemented by questionnaires sent to researchers and by records gathered from journals and online platforms. Most of the country cells were sampled.
- New Zealand: Volunteers visited more than 96% of the grid cells in both atlas editions. People were asked to indicate whether their survey was partial or complete in terms of habitat and species coverage when submitting their recording sheets to the atlas coordinators. The return of complete sheets varied across the country.
- New York: Volunteers selected a priority block to survey and were tasked with visiting each habitat within it. Paid workers were sent to remote areas to ensure adequate survey coverage. The goal was to record at least 76 species in every sampled block with breeding evidence reported for half of them.
- Alberta, Quebec, and The Maritimes: The data were collected across a 10×10 km grid in each study area. Volunteers were asked to spend ∼20 hours total surveying a given cell. However, the ∼20 hours of sampling served more as guidance than as a hard target, and variable survey durations are reported as sampling effort measures for each atlas.
- Czech Republic: Volunteers were required to cover all habitats within their assigned grid cell and record evidence of breeding for each observed species. The latest atlas also includes opportunistic records with identifiable breeding evidence from online portals and standardised 1-hour walks (timed-species-lists) in randomly chosen subsquares. The atlas covers most of the grid-cells in the country, except for a few remote locations along the country’s borders.
- United Kingdom (butterflies): Volunteers are encouraged to survey butterflies in all 10×10km grid cells over a 5-year period, as well as reporting opportunistic sightings. Almost all observations are reported at 1x1km grid resolution or finer. These data are combined with records from standardised butterfly monitoring (e.g. timed counts, fixed-route transect counts) into the Butterflies for the New Millennium recording scheme. The 5-year surveys run consecutively, and all records are verified by volunteer taxon experts. Only resident and regular migrant butterfly species are considered, while (re)introduced species and scarce migrant, vagrant and adventive species are excluded.
- Europe (EBBA): The data are compiled from country-level atlases, monitoring schemes, casual observations, and targeted surveys performed by the EBBA team. The data were standardised and assigned to a matching ∼50×50 km grid cell. National coordinators of data-providing countries were asked to evaluate the breeding categories for each species, adding an extra layer of data quality assessment.

After data collection, atlases have a vetting committee that evaluates the reliability of the records and seeks additional expert advice to ensure the final dataset is trustworthy. These semi-structured sampling procedures across a grid, accompanied by basic survey protocol guidance and post hoc evaluation, make gridded atlases suitable for pooling and comparison despite potential differences among them.

### Data filtering and cleaning

In the context of the planned follow-up analyses (not reported in this paper), we made the following decisions. The first atlas period for the Czech Republic (1973–1977) was not considered because the grid is not compatible with subsequent periods. Similarly, Japan’s most recent atlas period (2016–2021) was omitted given that the volume of voluntary reports was substantially higher and targeted different species relative to earlier survey events (which is not possible to correct using Frescalo). For all atlases, we retained only grid cells surveyed across all periods. This is because occupancy estimation relies on survey effort across periods, and a cell with zero effort in any one period would introduce an artefact into the occupancy calculation. For cells partially truncated by coastlines or administrative boundaries, we excluded those in which more than three-quarters of the cell area lay outside of the atlas survey area.

### From raw data to probabilities

We converted the raw atlas data (species detections) into occupancy probabilities using a two-step process. First, we estimated sampling effort for each grid cell within each atlas period, and then we ran a Bayesian spatial occupancy model for each species and atlas period, using the estimated sampling effort as the only predictor of detection in the observation process of the model. As the main output, we provide the estimated occupancy probability for each species across grid cells for each atlas period, along with the uncertainty around each estimate.

### Sampling effort calculation

Even though some of the atlases included information on the number of visits or hours spent recording in the grid cell, these values do not necessarily reflect sampling effort, as some recorders may be able to record more species than others within the same time span. Thus, to accurately estimate sampling effort across our datasets, we ran the Frescalo algorithm ^27^ for each atlas and atlas period using the R package sparta ^44^. This method uses the frequency of a set of “benchmark” species (i.e., species that are widespread and assumed to be consistently recorded), and compares their presence in a focal grid cell to their frequency in neighbouring cells, weighted by habitat similarity and physical distance. See Goury et al. ^45^ for a practical guide on the use of Frescalo.

We calculated the weights required to run Frescalo using the euclidean distances between grid cells and the mean elevation per cell (as a habitat similarity proxy), downloaded using *geodata* ^46^. These weights ensure that, within the neighbourhood of selected cells, species lists from cells that are physically more distant or less similar in elevation contribute less than those from nearer or more similar cells. We used 20 as the number of neighbours to include after ranking by distance, and 10 as the number of neighbours to include after ranking by similarity. Finally, we ran the algorithm and calculated the proportion of benchmark species in each grid cell and sampling period using the function *frescS_it*, developed by Auffret & Arnell ^47^. We used this proportion of benchmark species as our proxy for sampling effort.

### Occupancy models

To account for the effect of varying sampling effort in the raw data, we ran a spatial occupancy model for each species recorded in each atlas-period combination using the *spPGOcc* function from spOccupancy ^28^. This function fits a single-species spatial occupancy model using Nearest Neighbor Gaussian Processes (NNGP) and Pólya-Gamma data augmentation for fast and efficient computation (for more details, see Doser et al., 2022).

The model separates the observation process from the true occupancy. Thus, given that we have estimated the sampling effort per grid cell from the data (through Frescalo), we can use this variable to model the observation process, while assuming the true occupancy is the result of the observation process, given the area of the grid cell.

The occupancy of a species at a grid cell in a specific time period is therefore conditioned on the presence of the species, the area of the cell, and the estimated recording effort at that location. We did not include environmental covariates in the occupancy model for three reasons. First, identifying relevant predictors would require researching the habitat preferences of thousands of species individually. Second, using generic predictors (e.g., annual temperature, precipitation seasonality) would bias occupancy estimates toward species that respond strongly to those particular variables. Third, since we aim to test what drives occupancy change, including environmental predictors would introduce circularity into the analyses.

### Model description

Let *z*_*j*_be the true presence (1) or absence (0) of a species at a site *j*, with *j*=1, …, *j*. We assume this latent occurrence variable arises from a Bernoulli process following,

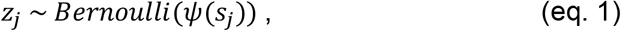

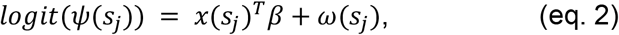

where ψ(*s*_*j*_) is the probability of occupancy at site *j* with coordinates *s*_*j*_, which is a function of site-specific covariates *x* and a vector of regression coefficients (β), T denotes transposition of column vector *x*(*s*_*j*_), and where ω(*s*_*j*_) is a realisation from a zero-mean spatial Gaussian process. In our case, the only site-specific occurrence covariate is the area of the grid cell, which accounts for the fact that, although the grids are almost regular, not all grid cells are exactly the same, some are cropped or elongated.

The Gaussian process *ω*(*s*_*j*_) introduces autocorrelation in the *ψ*(*s*_*j*_), based on the empirically well-supported assumption that most real-world species distributions are autocorrelated ^48,49^; this is also the reason why the model can separate the detection process (described below) from the occupancy *ψ*(*s*_*j*_), something that would otherwise require repeated visits. It is, however, also the reason the model has not converged for some of the extremely rare species (see the “Flagged species” section below).

The zero mean Gaussian process considered assumes that,

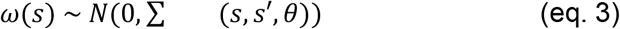

where 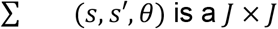 covariance matrix that is a function of the distances between any pair of site coordinates *s* and *s*′, and a set of parameters (*θ*) that govern the spatial process according to a spatial correlation function. We used the exponential function *θ* = {*σ*^2^, *ϕ*}, where *σ*^2^ is the spatial variance parameter and *ϕ* is a spatial decay parameter.

Now let *y*_*j*_ be the observed detection (1) or nondetection (0) of a species of interest at site *j*. We see the detection-no detection *y*_*j*_ data as arising from a Bernoulli observation process:

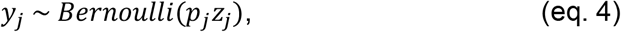

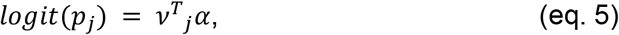

where *p*_*j*_ is the probability of detecting a species at site *j* (given it is present at site *j*), which is a function of site-specific covariates *v*, with *T* again denoting transposition of *v*, and a vector of regression coefficients (*α*). In our case, the only site-specific detection covariate is the sampling effort in the grid cell.

We assign multivariate normal priors for the occurrence (*β*) and detection (*α*) regression coefficients.

### Model fitting

We fitted the model using ‘exponential’ as the spatial correlation function and 5 nearest neighbours. We used 300,000 samples per chain (12,000 batches of length 25), burn-in 100,000, thinning rate 500, and 3 chains. The total posterior samples were 1,200 for each parameter and for each *ψ(s*_*j*_). These large numbers of iterations ensured convergence of the spatial covariance parameters. Finally, we stored the mean and the standard deviation (SD) of the occurrence probability of each species per grid cell and atlas period, among other parameters. See the *Data Records* section for a full description of the data.

The analyses were run in parallel on a server with two CPU AMD of 128 cores and 256 threads each and a memory of 550 GB. Using 150 cores, model fitting took on average 27.8 hours per atlas, depending on the number of species, grid cells and periods considered. In the case of the Czech Republic (233 species, 615 grid cells, and 3 periods), the models took around 11.4 hours, while on the other extreme, Europe (with 518 species, 2506 grid cells, and 2 periods) took 68.2 hours.

### Modelling performance

We checked model convergence through *Rhat* statistics and Bayesian p-values. *Rhat* compares within-chain and between-chain variance across multiple Markov Chain Monte Carlo (MCMC) chains ^50^, while the Bayesian p-value evaluates the proportion of model-generated data (from posterior samples) where the fit statistic is larger than the one from the real data ^51^. Specifically, we checked that the *Rhat* values were <1.1 for all estimated parameters (indicating that the chains have converged to the true posterior distribution), and that the Bayesian p-value was around 0.5, ranging from 0.1 to 0.9 (Doser et al. 2022). We also compared the raw occupancy of each species per grid cell *with* the estimated occupancy values and visually inspected the species’ probabilistic maps for clear outliers or distortions.

The *Rhat* values for all estimated parameters were mostly <1.1, except for the *σ*^2^ and *ϕ* parameters (for which convergence is always challenging in spatial occupancy models). Bayesian p-values ranged between 0.485 and 0.649 (mean = 0.566, SD = 0.05) across atlases. Based on the raw occupancy vs estimated occupancy values per grid cell and visual checks of the probabilistic maps of the species, the overall convergence rate per species across all periods ranged from 55.4% (New Zealand) to 100% (UK) (Table 2).

**Table 2.** Model performance. Total number of species modelled, number of species for which the spatial occupancy model performed poorly (at least in one atlas period), and percentage of species for which the occupancy model converged, for each atlas.

|  | CZ | NY | AL | JP | NZ | QE | TM | EU | UKB |
| --- | --- | --- | --- | --- | --- | --- | --- | --- | --- |
| # species modelled | 233 | 252 | 367 | 256 | 240 | 256 | 222 | 518 | 58 |
| # poor performing | 54 | 45 | 97 | 46 | 107 | 46 | 54 | 80 | 0 |
| % converged | 76.8 | 82.1 | 73.6 | 82 | 55.4 | 82 | 75.7 | 84.6 | 100 |

### Flagged species

We classified an occupancy value as inflated when the mean estimate (’mean.psi’) was at least 10 times the raw occupancy (i.e., the raw number of occupied grid cells divided by the total number of grid cells). In each atlas, several species were flagged for having inflated occupancy values (marked as “poor performing” in Table 2). This issue occurred only for rare species (e.g., with raw occupancy of 1-2% of the territory) that had very sparse spatial distributions, leading to inflated estimated occupancies. We advise users to interpret estimates for these species with caution. Additionally, we advise users to always take into account the occupancy (’mean.psi’) together with its uncertainty (’sd.psi’) and if possible, propagate the uncertainty in their analyses (e.g., Grattarola et al., ^52^).

These inflated estimates are closely tied to the challenges of modelling rare species. Assessing convergence in spatial factor models can be a complicated task ^53^. With so few observations, fitting any basic occupancy model is generally difficult, and thus a spatial occupancy model is even more challenging because there is limited information to estimate all parameters. The spatial model we use includes a site-level random effect that assumes occupancy is spatially correlated, meaning nearby sites tend to have similar occupancy probabilities (e.g., if a species occurs in one cell, it is more likely to occur in neighbouring cells than in distant ones). This is why a low number of records is less problematic when there is spatial autocorrelation in the detection pattern.

## Data Record

The data, including the occupancy probability for each species per grid cell within an atlas period and the uncertainty around each probability estimate, and the code used to generate them, are available at [anonymised repository: https://anonymous.4open.science/r/gridded-atlas-probabilities-2094/]. We provide a separate folder for each of the 9 atlases. See more details below.

### Data folder structure

Each folder is named after the dataset ID in Table 1, using the format *occ_<datasetID>*. For

example, “occ_5” corresponds to the Czech Republic dataset (datasetID = 5).

### Data files

Inside each folder, you will find three types of files: the species distribution probabilities (’occupancy_probabilities’), the diagnostics of all species in each period, including the flagged species (’diagnostics’), and the atlas grid (’atlas_grid’). The files are named according to the ID of the dataset, using the format *<file type>_<datasetID>*.*<file extension>*. For example, *occupancy_probabilities_5*.*duckdb* contains the data on species occupancy probabilities for the Czech Republic.

#### 1 ’*occupancy_probabilities’:* species grid-level data

This is a long table stored as a DuckDB database file (.*duckdb*), a self-contained, serverless database that organises data in a columnar layout optimised for efficient querying of large datasets without loading the entire table into memory ^54^. The file contains the occupancy probability data for a species during a given atlas period, and includes the mean occupancy probability and its standard deviation for each grid cell (’mean.psi’, ’sd.psi’). Each element of the list is named using the scientific name of the species (’sp.name’), which matches the same value in the ’diagnostic’ object.

DuckDB database file with one row per grid cell and the following variables:

- period: Integer that refers to the period number (e.g., 1,2,3)
- sp.name: Standardised scientific name of the species
- siteID: Unique identifier of the grid cell (maps to siteID in ’atlas_grid’)
- mean.psi: Mean occurrence probability (*ψ* ∈ [0, 1])
- sd.psi: Standard deviation of the occurrence probability
- mean.w: Mean spatial random effect
- sd.w: Standard deviation of the spatial random effect
- mean.p: Mean detection probability (*p ∈* [0, 1])
- sd.p: Standard deviation of the detection probability

#### 2 ’*diagnostics’:* species-level model parameters and performance diagnostics

This CSV file (.csv) contains species-level model parameters and estimates of model performance. Species were flagged (’is.flagged’ = TRUE) if, in any atlas period, ’mean.psi’ was at least ten times greater than the raw occupancy (the proportion of occupied grid cells out of the total number of grid cells) in any atlas period.

CSV file with one row per species and the following variables:

- sp.name: Standardised scientific name of the species
- alpha.det: Mean detection regression coefficient (*α*)
- beta.occ: Mean occurrence regression coefficient (*β*)
- bayesian.p: Bayesian p-value from posterior predictive checks (i.e., the proportion of posterior samples in which the fit statistic of the model-generated data exceeds that of the observed data; values near 0.5 indicate a good fit)
- rhat.beta.int: Diagnostic statistic for the occurrence intercept
- rhat.beta.area: Diagnostic statistic for the occurrence coefficient for area (*β*)
- rhat.alpha.int: Diagnostic statistic for the detection intercept
- rhat.alpha.eff: Diagnostic statistic for the detection coefficient for effort (*α*)
- rhat.sigma: Diagnostic statistic for the spatial variance (*σ*^2^)
- rhat.phi: Diagnostic statistic for the spatial range (*ϕ*)
- is.flagged: Logical (TRUE/FALSE), indicating whether the species has a poorly performing model.

#### *’atlas_grid’:* spatial grid for the atlas

This is a geopackage file (.gpkg), a self-contained, standards-based format built on SQLite that stores vector geometries, attribute tables, and spatial reference information in a single portable file without requiring dedicated GIS software or a database server. The file stores the spatial polygons in WGS84 (World Geodetic System 1984) of the entire atlas grid. It provides the geometric reference to which the ’occupancy_probabilities’ estimates are spatially linked via siteID.

Geopackage file (POLYGON geometry) with one row per grid cell and the following variables:

- siteID: Unique identifier of the grid cell (maps to siteID in ’occupancy’)
- datasetID: Identifier of the dataset
- footprintSRS: Spatial reference system of the grid cell footprint
- verbatimFootprintSRS: Original spatial reference system as provided in the source data
- area: Total area of the grid cell (km^2^)
- croppedArea: Area of the grid cell after cropping to the study region boundary (km^2^)
- areaUnit: Unit of area measurement
- croppedAreaPercent: Proportion of the grid cell area retained after cropping (%)
- geometry: Polygon geometry of the grid cell

### Data Overview

To illustrate our analyses, we present the full data generation process and results using the New York state atlas as a case example. See the Code Availability section for more details.

The Frescalo run took 2 minutes. We checked whether the proportion of benchmark species calculated were comparable to the raw numbers of effort (on the original data) (Figure 2ab). We found high levels of correlation between raw and estimated sampling effort metrics (*τ* = 0.18, p < 0.001; *ρ* = 0.257, p < 0.001; Figure 2c). These metrics are not expected to correlate perfectly, since the same number of visits across grid cells can still reflect very different levels of sampling effort ^27^. Nonetheless, the overall pattern aligns as expected: poorly sampled grid cells tend to show a lower proportion of benchmark species (Figure 2c).

**Figure 2.**
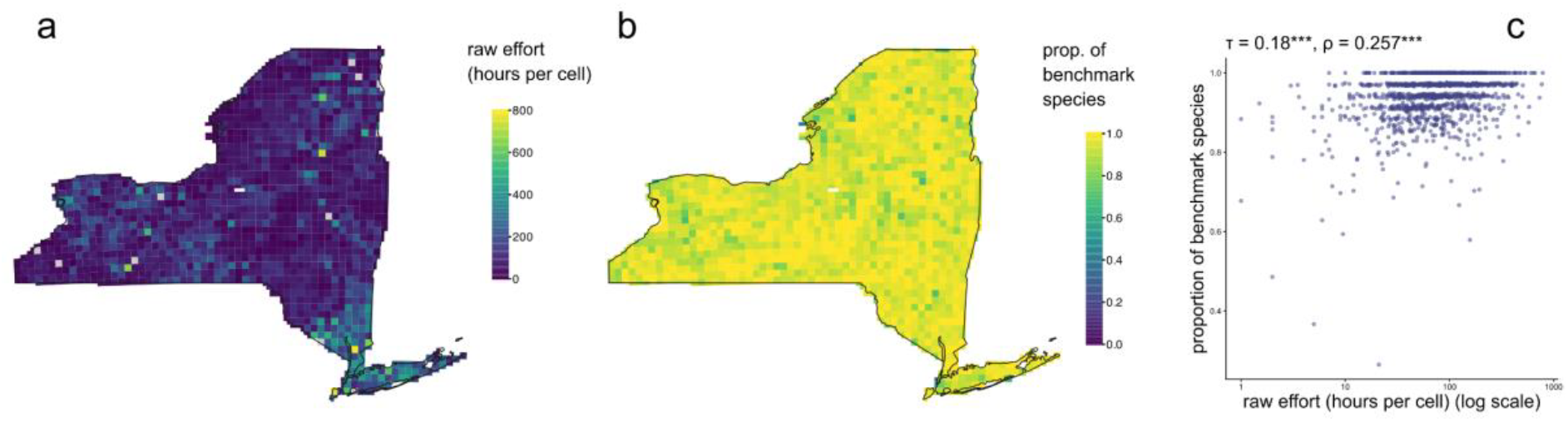
Association between the raw effort (number of visits as in the original data) and the proportion of local benchmark species per grid cell, using the first edition of the New York state atlas (1980-1985) as an example. (a) Distribution of the raw effort (number of hours per cell) (log scale), (b) distribution of the proportion of benchmark species (calculated using Frescalo), and (c) correlation between raw sampling effort (hours per cell) and estimated sampling effort per grid cell (proportion of benchmark species). Kendall’s τ and Spearman’s ρ correlation values are shown on top of the panel. The *** symbols indicate *p value* < 0.001.

The occupancy model took ∼19 hours to run (252 species, over 1,363 grid cells, and atlas 2 periods). We flagged 45 species as poor performing (17.9% of the total). We performed visual checks of the probabilistic maps of the species, and present *Setophaga citrina* as an example (Figure 3).

**Figure 3.**
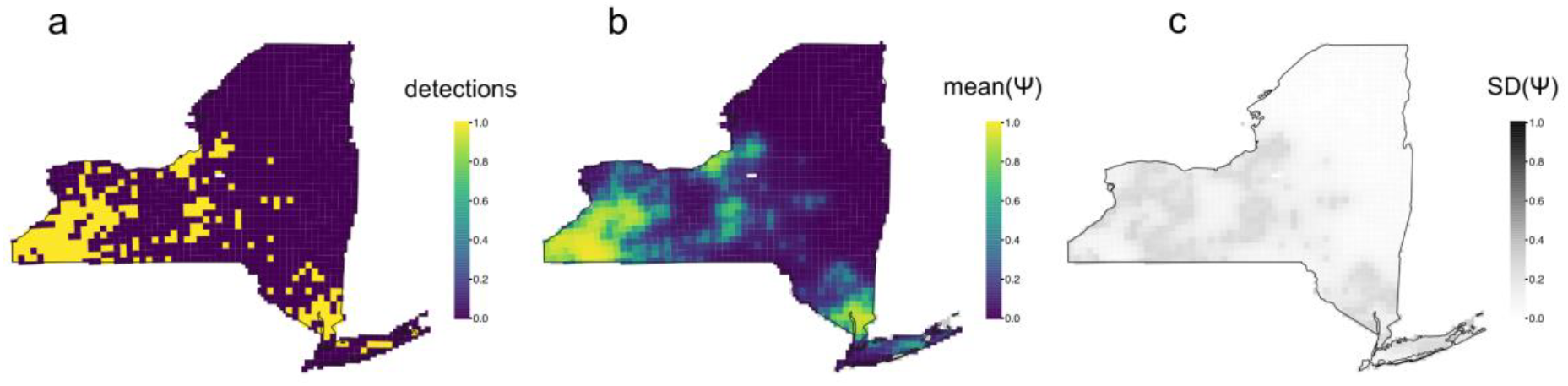
Example of estimated occupancy probability using *Setophaga citrina* from the first edition of the New York state (1980-1985) as an example. (a) Raw occupancy (detections) per grid cell, (b) mean occupancy probability ψ per grid cell, and (c) uncertainty (SD) of the occupancy probability per grid cell.

### Usage Notes

The data are available under the CC-BY license (https://creativecommons.org/licenses/by/4.0); reusers of the data must acknowledge the authors but are allowed to distribute, remix, adapt, and build upon the created material. For inquiries about the raw data and their accessibility, and for any further collaborations, we encourage reusers to contact each atlas’ point of contact (Table 1).

To help users access and process the data, we provide an html vignette that demonstrates how to: filter atlases, atlas periods, and species (including flagged ones), and obtain their distribution probabilities to create species maps. We recommend using R packages ‘sf’ ^55^, ‘duckdb’ ^54^, and ‘tidyverse’ ^56^ to manipulate the data. Importantly, the data have been made fit-for-purpose (with standardised species name and geometries), and thus do not need to be cleaned any further.

## Data Availability

The database can be accessed and downloaded at https://anonymous.4open.science/r/gridded-atlas-probabilities-2094/, under a CC-BY license (https://creativecommons.org/licenses/by/4.0).

## Code Availability

The code to produce the occupancy data is available at https://anonymous.4open.science/r/gridded-atlas-probabilities-2094/, under a GNU General Public License license (https://www.gnu.org/licenses/gpl-3.0.html).

The code for each function is annotated, stating the purpose, inputs, outputs, and package dependencies needed. To demonstrate how to run the Frescalo algorithm and the spatial occupancy model, we provide a vignette with a case example using data from the state of New York (see *Data Overview*). All atlases were processed following the same method.

## Acknowledgements

Thanks to Jeff Doser for important comments related to spOccupancy and spatial occupancy models. Bird atlas data are mostly collected by skilled volunteers coordinated by ornithological organisations, which are responsible for the quality of the observations, data curation, analyses, and publication. We wish to acknowledge all these fieldworkers and the organisations that made these atlas possible, i.e. Czech Society for Ornithology, New York State Breeding Bird Atlas, Nature Alberta, Environment Agency of Japan and Biodiversity Center of Japan, Ornithological Society of New Zealand, Québec Breeding Bird Atlas, Maritimes Breeding Bird Atlas, the European Bird Census Council, and Butterfly Conservation and Natural England, as well as all their partners and supporters. FG, GOS, MT, CDS, FJRW, and PK were funded by the European Union (ERC, BEAST, 101044740). LH was funded by the Heather Corrie Fund by Butterfly Conservation.

## Author Contributions

Conceptualisation and Investigation: FG and PK. Resources and Data curation: FG, GOS, MT, CDS, FJRW, RF, LH, SH, and PK. Formal analysis, Methodology, Visualisation, and Writing – original draft: FG. Funding acquisition: PK. Writing – review & editing: all authors.

